# Enter the matrix: factorization uncovers knowledge from omics Names/Affiliations

**DOI:** 10.1101/196915

**Authors:** Genevieve L. Stein-O’Brien, Raman Arora, Aedin C. Culhane, Alexander V. Favorov, Lana X. Garmire, Casey S. Greene, Loyal A. Goff, Yifeng Li, Aloune Ngom, Michael F. Ochs, Yanxun Xu, Elana J. Fertig

## Abstract

Omics data contains signal from the molecular, physical, and kinetic inter- and intra-cellular interactions that control biological systems. Matrix factorization techniques can reveal low-dimensional structure from high-dimensional data that reflect these interactions. These techniques can uncover new biological knowledge from diverse high-throughput omics data in topics ranging from pathway discovery to time course analysis. We review exemplary applications of matrix factorization for systems-level analyses. We discuss appropriate application of these methods, their limitations, and focus on analysis of results to facilitate optimal biological interpretation. The inference of biologically relevant features with matrix factorization enables discovery from high-throughput data beyond the limits of current biological knowledge—answering questions from high-dimensional data that we have not yet thought to ask.

## Determining the dimensions of biology from omics data

High-throughput technologies ushered in an era of big data in biology [1,2] and empowered *in silico* experimentation which is poised to characterize **complex biological processes (CBPs)** (RNAseq example given in Fig 1)[3]. The natural representation of high-dimensional biological data is a matrix of the measured values (expression counts, methylation levels, protein concentrations, etc.) in rows and individual samples in columns (Fig 2, Key Figure). Columns corresponding to experimental replicates, or samples with similar phenotypes will have values from the same distribution of biological variation. The related structure in the data is observed because they share one or more CBP. The activity of CBPs need not be identical in each sample. In these cases, the values of all molecular components that are associated with a CBP will change proportionally to the relative activity of that CBP. These phenotypes and CBP activities are often unknown a priori, requiring unsupervised computational techniques to learn them directly from the biological data.

**Figure 1.**
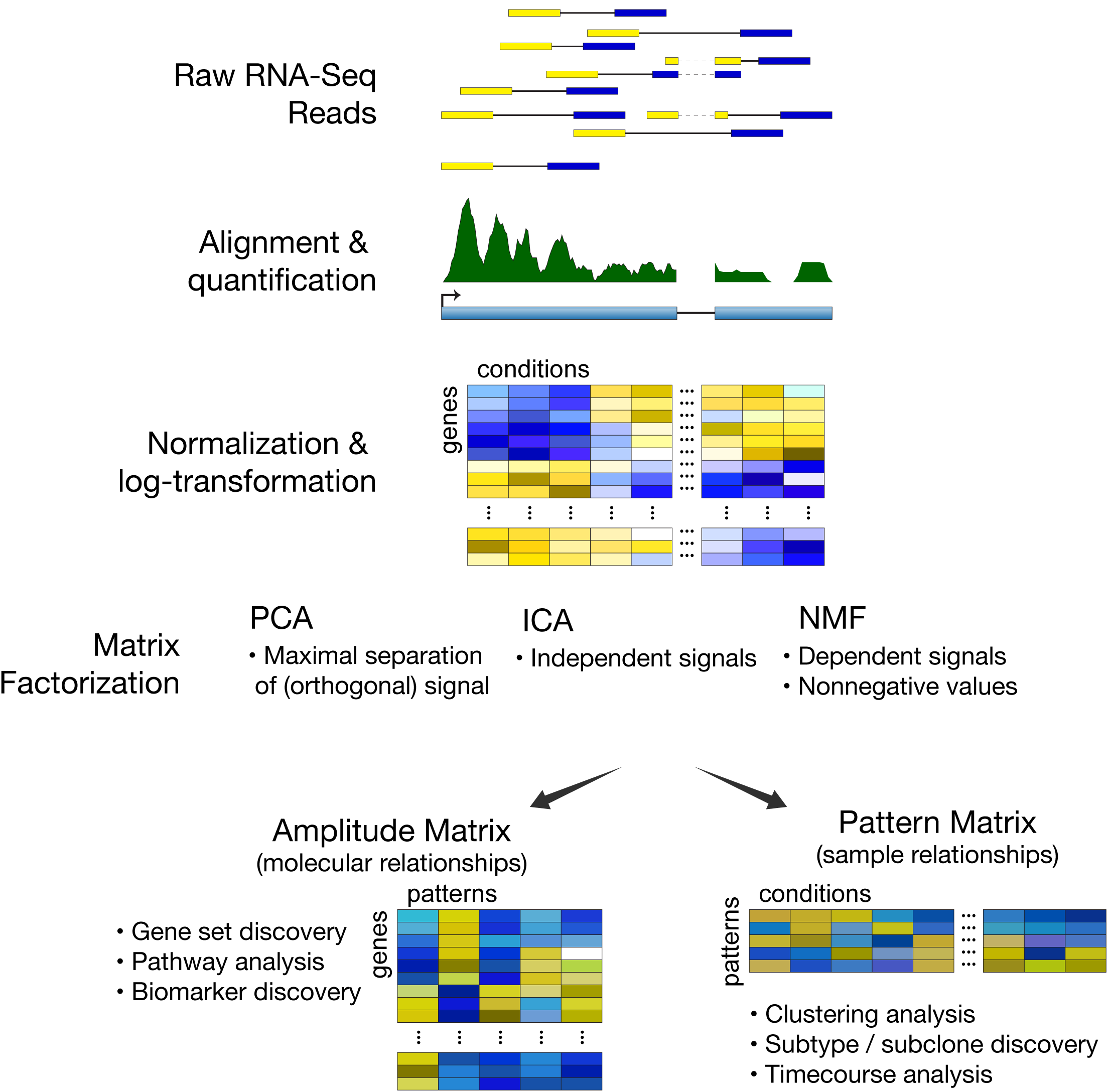
Pipeline for matrix factorization analysis of RNA-seq data. First, the RNA-seq data is preprocessed by alignment, gene-level quantification, and normalization. The data must aslo be log transformed for most matrix factorization methods, with the exception of specialized methods that have been developed for count data. Three dominant types of MF problems can be applied to analyze this data: Principle Component Analysis (PCA), Non-negative Matrix Factorization (NMF), or Independent Component Analysis (ICA). Each of these problems find distinct molecular and sample relationships in the amplitude and pattern matrices, respectively. The Amplitude matrix learned that reflects molecular relationships can be used for gene set analysis, pathway analysis, and biomarker discovery. The Pattern matrix that reflects sample relationships can be used for clustering, subtype / subclone discovery, and timecourse analysis.

**Figure 2.**
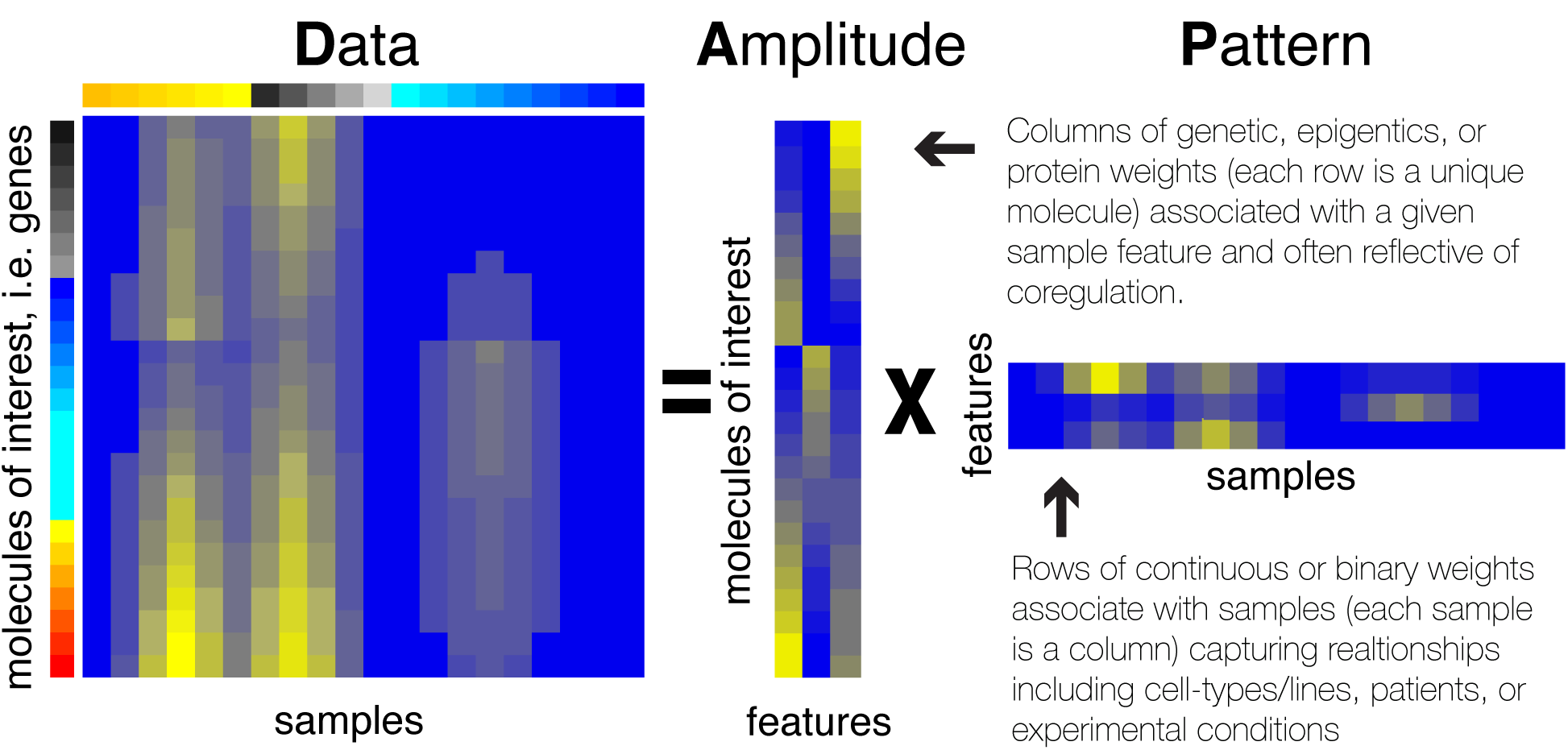
Omics technologies yield a data matrix that has each sample as a column and each observed molecular value (expression counts, methylation levels, protein concentrations, etc.) as a row. This data matrix is preprocessed with techniques specific to each measurement technology and then input to an MF technique for analysis. MF decomposes the preprocessed data matrix into two related matrices that represent its sources of variation: an amplitude matrix and a pattern matrix. The rows of the amplitude matrix quantify the sources of variation among the molecular observations and the columns of the pattern matrix quantify the sources of variation among the samples. The matrix product of the amplitude and pattern matrices approximates the preprocessed input data matrix. The number of columns of the amplitude matrix equals the number of rows in the pattern matrix, and represents the number of dimensions in the low-dimensional representation of the data. Ideally, a pair of one column in the amplitude matrix and the corresponding row of the pattern matrix represents a distinct source of biological, experimental, and technical variation in each sample (called complex biological processes, CBPs). The values in the column of the amplitude matrix then represent the relative weight of each molecule in the CBP and the values in the row of the pattern matrix its relative role in each sample.

The relationships between CBPs and similarities between samples constrain high-dimensional datasets to have low-dimensional structure. The number of genes, proteins, and pathways that are concurrently active within any cell is constrained by its energy and free-molecule limitations [4]. Only a characteristic subset of CBP will be active in any cell at a given time. Thus for a dataset where columns share CBP, a low dimensional structure can be extracted which is smaller than either the number of rows or the number of columns. Matrix Factorization (MF) is a class of unsupervised techniques provide a set of principled approaches to parsimoniously reveal the low-dimensional structure while preserving as much information as possible from the original data (see Box 1).

When applied to high-throughput omics data, MF techniques learn two matrices: one describes the structure between features (e.g., genes) and another the structure between samples (Fig 2). Here, we call the former gene-level matrix the **Amplitude matrix** and the latter sample-level matrix the **Pattern matrix**. There are numerous approaches to MF, including both gradient-based and probabilistic methods (Box 1). Additional terms have been coined for the Amplitude and Pattern matrices based upon the MF problem applied and on the specific application to high-throughput biological data. Other reviews discuss the mathematical and technical details of MF techniques [5–8] and their applications to microarray data [9].

Here, we focus on the biological applications of MF techniques and the interpretation of their results since the advent of sequencing technologies. We describe a variety of MF techniques applied to high-throughput data analysis and compare and contrast their use for biological inference. Many techniques described are for sequencing data that is preprocessed with log transformation [10] or models of sequencing depth [11], while others directly model read counts [12]. We focus on examples in pathway analysis, subtype and clonal identification, time course analysis, multi-omics integration, and single cell data to present a field with much wider applications.

## Data-driven gene sets from MF provide context-dependent coregulated gene modules and pathway annotations

Genomics data are often interpreted by identifying molecular changes in sets of genes annotated to functionally related modules or pathways, called gene sets [13,14]. Often the association between gene sets and functions used are based upon human curation of the literature [15,16]. Such set-level interpretations often lack important contextual information [13,17,18] and cannot describe genes with unknown function or genes associated with new functional mechanisms.

The amplitude matrix from MF analysis can be used both for literature-based gene set analysis and to define new data-driven gene signatures (Fig 3). The values in each column of these amplitude matrices are continuous weights describing relative contribution of a molecule in each inferred factor. In cases where factors distinguish CBPs, the relative weights of these molecules can be associated with functional pathways. The same molecule may have high values in multiple columns of the amplitude matrix. Thus, MF techniques are able to account for the cumulative effect of genes that participate in multiple pathways. The properties of the amplitude matrix, and subsequently the interpretation of their values, depend critically on the specific MF problem and algorithm selected for analysis.

**Figure 3.**
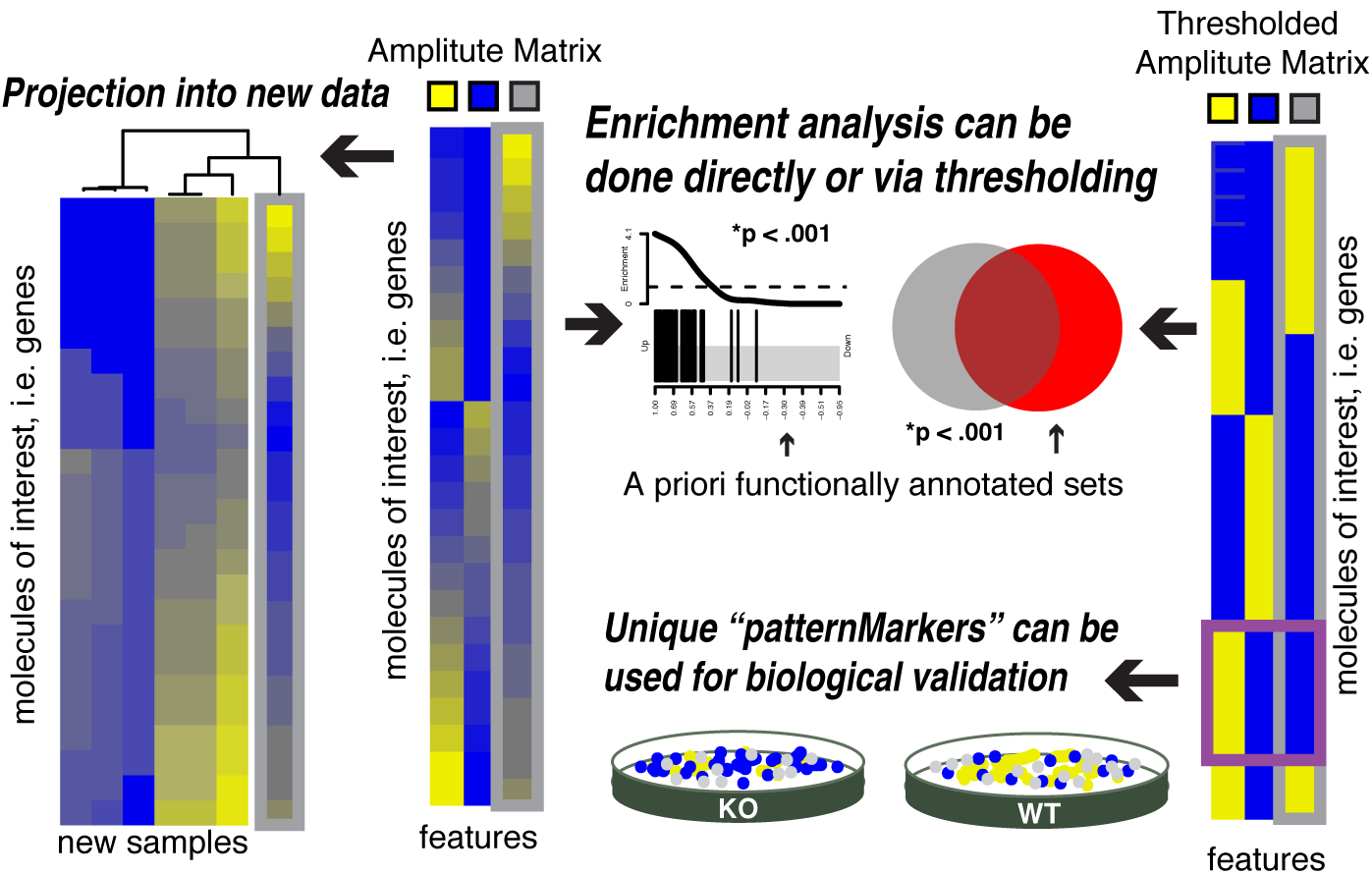
The amplitude matrix from MF can be used to derive data-driven molecular signatures associated with a CBP. The columns of the amplitude matrix contain continuous weights describing relative contribution of a molecule in a CBP (middle). The resulting molecular signature can be analyzed in a new dataset to determine the samples in which each previously detected CBP occurs, and thereby assess function in a new experiment. This comparison may be done by comparing the continuous weights in each column of the amplitude matrix directly to the new dataset (left). The amplitude matrix may also be used in traditional gene set analysis (right). Traditional gene set analysis using literature curated gene sets can be performed on the values in each column of the amplitude matrix to identify whether a CBP is occurring in the input data. Data-driven gene sets can also be defined from this matrix directly using binarization, and used in place of literature curated gene sets to query CBPs in a new dataset. Sets defined from molecules with high-weights in the amplitude matrix comprise signatures akin to many curated gene set resources whereas molecules that are most uniquely associated with a specific factor (purple box) may be biomarkers.

The three most prominent MF approaches are **Principal Component Analysis (PCA), Independent Component Analysis (ICA)**, and **Non-negative Matrix Factorization (NMF)**. Each of these techniques has a distinct mathematical formulation of a distinct MF problem that is described in the Box 2 and other reviews [5,8,19–22]. Briefly, PCA finds dominant sources of variation in high-dimensional datasets, inferring genes that distinguish samples. Maximizing the variability captured in certain factors, as opposed to spreading relatively evenly among factors, may mix the signal from multiple CBPs in a single component. Therefore, PCA may conflate processes that sometimes occur and make interpretation of the amplitude matrix for define data-driven gene sets difficult.

To learn distinct processes, ICA learns factors that are statistically independent, resulting in more accurate associate with literature-derived gene sets [23–25]. Comparison analyses in Rotival et al [26] found that ICA could identify modules with known biological function. NMF methods constrain all elements of the amplitude and pattern matrices to be greater than or equal to zero [27,28]. NMF is well suited to transcriptional data, which is typically non-negative itself, and semi-NMF is also applicable to data that can have negative values. The assumptions of NMF model both the additive nature of CBPs and parsimony, generating solutions that are biologically intuitive to interpret [29].

The solutions from both ICA and NMF may vary depending upon the initialization of the algorithm, leading to disparate amplitude matrices. Therefore, it is critical to ensure that particular solution used for analysis provides an optimal and robust solution before using the amplitude matrix to define data-driven gene signatures. Bayesian techniques to solve NMF were found to have more robust amplitude matrices than gradient-based techniques, and thus more accurate associations of the values in the amplitude matrix with functional pathways [5,30]. These associations also depend critically on the input data. Therefore, to learn context dependent genes sets, MF can be applied to datasets with well-defined experimental perturbations [31].

Standard gene set analysis can be applied directly to the values in each column of the amplitude matrix to associate the inferred factors with literature-curated sets. New, context dependent gene sets can also be learned from the values in the amplitude matrix. Gene set annotations are often binary. Thresholding techniques to select which genes belong to a pathway from the amplitude matrix for binary membership provide an output similar to gene sets in databases [31,32]. Other studies also integrate the literature-derived gene signatures in these thresholds to refine the context of pathway databases [30,33]. The genes derived from these binarizations can be used as inputs to pathway analyses from differential expression statistics in independent datasets (Fig 3, right) and are analogous to the hierarchical-clustering based gene modules [34] and gene expression signatures from public domain studies in the MSigDB gene set database [35]. Another means of binarizing the data is to find genes that are most uniquely associated with a specific pattern to use as biomarkers of the cell type or process associated with that pattern [36,37]. Selecting genes based upon these statistics can facilitate visualization of the CBPs in high-dimensional data [36]. Whereas binarization of genes with high weights can associate a single gene with multiple CBPs, the statistics for unique associations link a gene with only one CBP. Therefore, these statistics also define specific genes that may be biomarkers of the cell type/state or a process [36] (Fig 3, right).

Although binary pathway models are substantially easier to interpret, continuous values from the original factorization provide a better model of the input data. Weighted gene signatures have been shown to be more robust to noise and missing values in the data [38]. If a gene’s expression level is poorly measured in a sample, other genes in the same factor can imply the actual expression level of the gene in question. By considering each gene in the context of all other genes, factorization improves the robustness of findings. Further, continuous signatures can be associated directly with other samples using projection methods [38,39] or profile correspondence methods [40].

MF has also been applied to learn functional gene modules on non-transcriptional datasets. Alexandrov et al applied NMF to the mutational landscape of tumors defined from the number of nucleotides that differ from the reference genome in the context of the preceding and following nucleotides [41]. This analysis defined mutational signatures associated with distinct cancer driving processes [42]. Additional extensions have been applied to learn allele combinations in phenotypes [43], transcript-regulation of genes [44], distributions of transcript lengths [45], and discriminate peptides in mass spectrometry proteomics [46].

## MF learns relationships between samples that represent population stratification, tissue composition, cell types, disease subtypes, and clonality

Whereas each column of the amplitude matrix describes the relative contribution of molecules to a factor, each row of the pattern matrix describes the relative contribution of samples to a factor (Fig 2, Fig 4). Sample groups can be learned by comparing the relative weights in each row of the pattern matrix (Fig 4). The pattern matrix from MF can also be binarized to perform clustering [47,48] or kept as continuous values to define relationships between samples [49–51]. Just as molecules with high weights within a column of the amplitude matrix are associated with a common pathway, samples with high weights within a row of the pattern matrix can be assumed to share a common phenotype or CBP. Although here we refer to each of these sample-level matrices as the Pattern matrix, numerous other terms have been adapted based on the MF method and its application (Box 3).

**Figure 4.**
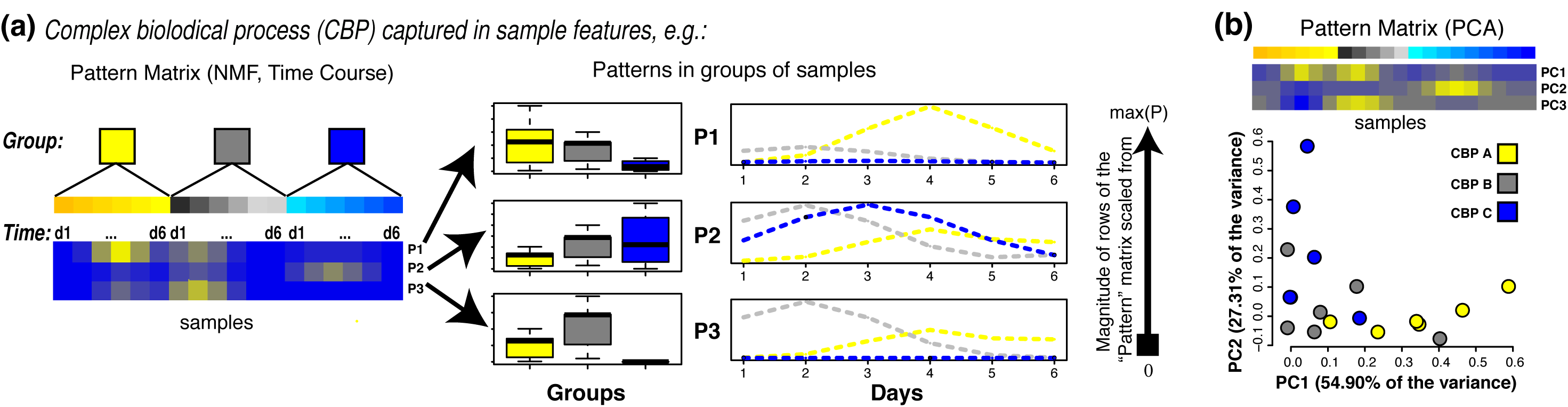
The pattern matrix from MF describes sample participation in a CBP. Trends or groupings of samples in a row of the pattern matrix can be tested against additional sample metadata to further define the biological details of a given CBP. Depending on the MF method used different visualizations can highlight specific aspects of the biology. **A.** Nonnegative matrix factorization method find the relative values of correlated or dependent measures just as pathway usage or time course data. If sample grouping are known *a priori* the relative usage of a CBP can then be inferred by comparing the weights of the pattern matrix between groups. Similarly, if samples correspond to time points the rows of the pattern matrix can be plotted as a function of time and sample condition to infer the dynamics of CBPs. **B.** Principal component analysis (PCA) on the other hand, uses an orthogonality constraint to maximize the amount of variance captured in each principal components (PC). Thus, it is often informative to plot samples projected into the first two PCs (right) to assess sample clustering.

The application of PCA to SNP data from 3,000 European individuals [52] demonstrates inference of sample co-relationships using the pattern matrix and found that much of the variation in DNA sequence is explained by the longitude and latitude of an individual’s country of origin. Additionally, statistical models can be formulated assuming that inheritance of an individual arises from proportions of ancestry in distinct populations through genetic admixture [52]. A MF based technique called sparse factor analysis also distinguishes these populations using GWAS data [53]. These analyses demonstrate that the ancestry of each individual is a dominant source variation in DNA sequence. Additional sources of variation in GWAS data arise from variants that give rise to disease risk, which can be shared among individuals with diverse genetic backgrounds [54]. These different sources of signal can give different sample groupings that reflect the biology of population genetics. For example, biologically a grouping inferred from GWAS which distinguishes ancestry is equally valid to a grouping inferred from the same GWAS data which distinguishes disease risk.

A single factorization of complex datasets can find multiple distinct sources of variation. For example, the power of MF to identify multiple sources of variation was seen when multiple technical factors from sample processing and biological factors were discovered in an ICA of gene expression profiles of 198 bladder cancer samples [55]. One factor in the pattern matrix of this analysis defined a CBP associated with gender. Because ICA simultaneously accounts for multiple factors in the data as separate rows in the same matrix, each row can fully distinguish a single biological grouping from the data.

Applying multiple types of MF techniques to the same dataset or a single MF to distinct subsets of a dataset can also find distinct sources of variability. For example, an NMF-class algorithm separated tissue-specific patterns from gene expression data for postmortem samples in the Genotype-Tissue Expression (GTEx) project [56]. This algorithm found a pattern that combined all samples from brain regions when applied to all tissue samples in GTEx, but separated the distinct brain regions when applied to only tissue samples from the brain. A different sparse NMF algorithm called CoGAPS simultaneously separated these brain regions from GTEx with additional patterns that are associated with the individual who donated those samples [36,57]. Both of these algorithms are equally valid, and their distinct formulation gives rise to the distinct features observed in the data. Applications of multiple types of MF techniques, or even the same MF algorithm with different parameters, may infer several CBPs or phenotypes within a single dataset, in essence providing answers to different questions.

Analysis of a single dataset with one MF algorithm using different numbers of factors can reflect a hierarchy of biological processes. For example, applying CoGAPS to data from a set of head and neck tumors and controls for a range of dimensionalities was able to separate tumor and normal samples when limited to two patterns, but further decomposed the tumor samples into the two dominant clinical subtypes of head and neck cancer when identifying five patterns for the same data [58]. The hierarchical relationship between patterns has been used to assess the robustness of patterns to quantify the optimality of the factorization [59] and learn the optimal dimensionality of the factorization [60]. Other algorithms use statistical metrics to estimate the number of factors [12,61]. While these algorithms quantify fit to the data, they may disregard the hierarchical nature of distinct CBPs learned by factoring biological data into multiple dimensions. This observation highlights the complexity of estimating the number of factors for optimal MF analysis of biological data (see Outstanding Questions).

Intra- and inter-tumor heterogeneity introduce a further degree of complexity to MF analysis of biological variation in their molecular data. **Computational microdissection** algorithms estimate the proportion of distinct cell types within a bulk samples by applying MF to genes whose expression are uniquely associated with each cell type [62]. Subsetting the data to different genes may give rise to different factors that represent different CBPs. Nonetheless, CoGAPS analysis of data subsets that were obtained by selecting equally sized sets of random genes found that the pattern matrices were consistent for each random gene set in expression data[36,63]. These results suggest that the dependency of a MF on the specific genes used for analysis may depend on the heterogeneity of the signal in the data matrix.

Even a pure tumor tissue can contain numerous subclones due to the accumulation of different driver events during tumor evolution. New MF techniques have been developed to estimate the proportion of the tumor that arises from each subclone [61,64–67]. Assumptions about the evolutionary mechanisms of the accumulation of molecular alterations can also be encoded in the factorization to model the resulting heterogeneity of these clones [12,61]. These studies demonstrate that encoding prior knowledge into MF can focus the resulting factors to reflect one of the equally valid biological groupings within the data.

## From snapshots to moving pictures: simplifying time course analysis

Entwined in the challenge of decomposing cell types and subpopulations is the fact that CBPs change over time. High-throughput time course datasets are emerging in the literature to account for the dynamics of biological systems. The central goal of time course analysis is to determine the extent to which molecules change over time in response to perturbations (e.g. developmental time, environmental factors, disease processes, or therapeutic treatments). Associating molecular alterations often relies on specialized bioinformatics techniques for time course analysis [68,69]. MF analyses can naturally infer changes in CBPs over time when applied to time course data because the continuous weights for each sample in the pattern matrix can vary across samples collected across distinct time points. The relative weights of rows of the pattern matrix can encode the timing of regulatory dynamics directly from the data (Fig 4A).

Both ICA and NMF were found to have signatures characterizing the yeast cell cycle and metabolism in early time-course microarray experiments [70,71]. The Sparse NMF techniques using Bayesian method had patterns that reflected the smooth dynamics of these phases [30,70]. This approach has been shown to simultaneously learn pathway inhibition and transitory response to chemical perturbation of cancer cells [72] and relate the the changes in phospho-proteomics trajectories between multiple therapies [73]. Similar analysis of healthy brain tissues learned the dynamics of transcriptional alterations common to the ageing process from multiple individuals [63]. MF techniques designed for cancer subclones described in the previous section have also been applied to repeat samples to learn the dynamics of cancer development, elucidating the molecular mechanisms that give rise to therapeutic resistance and metastasis. Even if the same number of biological features exist, the rate or timing of related features in different molecular modalities may be offset [74]. These discrepancies by data modality suggest that different regulatory mechanisms may be responsible for initiating and stabilizing the malignant phenotype [74].

## Integrated analysis of multiple omics data

Multi-omics data are generated in order to elucidate the molecular networks that govern phenotypes. MF can be applied to learn shared features between datasets [7,8,75]. Integration may occur between datasets with distinct samples measured with the same molecular type using different measurement technologies or in distinct technical batches. For example, an analysis of multiple microarray studies of the same cell lines across different platforms with an MF approach designed to find components that maximize covariance or correlated information discovers which microarray platforms have the most informative set of genes [76]. Gene regulation can also be inferred by applying MF to data with different molecular components of the same samples. For example, repression of gene expression by promoter methylation had been encoded to integrate gene expression and DNA methylation data of tumor samples [58]. Other techniques have used joint modeling of features learned separately in different data types to perform integrated inference [77–79]. Techniques that extend this integrated MF framework, including Bayesian group factor analysis [8] and tensor decomposition [80] can also identify common and specific factors among different molecular levels [81]. Developing such data integration techniques is an active area of research in both genomics and computational sciences. When mature, these techniques will be able to formulate complete gene regulatory networks to further systems biology.

## MF enables unbiased exploration of single cell data for phenotypes and molecular processes

MF approaches are a natural choice in Single-cell RNA-sequencing (scRNA-seq) data analysis due to its high dimensionality and are used to identify and remove batch effects, summarize CBPs, and annotate cell types in the data [82–85]. Whereas the analysis in bulk data dissects groups from a small subset of samples, the analysis in scRNA-seq data aggregates cells into groups of common cell types or CBPs [37,86]. Often these analyses are performed on a subset of the data containing the most variable genes. Newer computationally efficient methods are being developed to enable factorization of large omics datasets for genome-wide analysis [79]. Biological knowledge can be encoded with a class of MF algorithms that summarizes factors using gene sets [82,87,88].

Most MF techniques developed for bulk omics data are linear, namely they assume the gene expression changes from CBPs are additive. This assumption is violated in scRNA-seq data. One reason for the violation of the linearity assumption in MF is the the inability to distinguish true zeros from missing values. Imputation methods for preprocessing [84,89] or newer MF algorithms that model missing data are essential for scRNA-seq data. Branching of trajectories of cellular states and lack of synchronization of cell cycle in scRNA-seq data further violate the linearity assumption in MF. New non-linear factorization techniques are being developed to enhance visualization of trajectory structures in single cell data [83,90–92] in these cases.

## Concluding remarks

MF is a versatile class of techniques with broad applications to unsupervised clustering, biological pattern discovery, component identification, and prediction. Since MF was first applied to microarray data analysis in the early 2000s [70,93–95], the breadth of MF problems and algorithms for high-throughput biology has grown with their broad applications. MF problems are ubiquitous in the computational sciences, with examples including unsupervised feature learning [96–101] clustering and metric learning [102–104], subspace learning [105–110], multiview learning [111], matrix completion [112], multi-task learning [113] semi-supervised learning [114], compressed sensing [115], and similarity-based learning [116,117]. Dimension reduction of biological data with MF highlights perspectives and questions that investigators have not yet considered, and also enables tractable exploration of otherwise massive datasets.

Different classes of techniques solve MF, including gradient-based and probabilistic methods (see Box 1). Distinct MF problems each aim to identify certain types of features, in some cases different algorithms will learn distinct features from the same dataset. Therefore, investigators may benefit from applying multiple techniques with different properties, or by carefully considering the dataset and question to select exactly the right technique for that question. The features MF techniques extract are constrained by the dataset used to train them. These algorithms cannot learn unmeasured features nor can they correct for complete overlap between technical artifacts and biological conditions. Thus, being mindful of experimental design when selecting datasets and choosing those that are broad enough to cover the relevant sources of variability are essential. Advances to MF and related techniques will be essential to powering systems level analyses from big data (see Outstanding Questions).

## Acknowledgements

We thank Orly Alter, J Brian Byrd, Michael Love, Irene Gallego Romero, Lillian Fritz-Laylin, Luciane Kagohara, Louise Klein, Craig Mak, Matthew Stephens, Daniela Witten, and other members of New PI Slack for their insightful feedback.

This work was supported by National Institutes of Health [grant numbers NCI 2P30CA006516-52 and 2P50CA101942-11 to A.C.C., NCI R01CA177669 E.J.F., NLM R01LM011000 to M.F.O. and NCI P30 CA006973], the Johns Hopkins University Catalyst and Discovery Awards to E.J.F., the Johns Hopkins University IDIES Award to E.J.F. and RA, the Johns Hopkins School of Medicine Synergy award to E.J.F. and L.A.G., a grant from The Gordon and Betty Moore Foundation (GBMF 4552) to CSG, Alex’s Lemonade Stand Foundation’s Childhood Cancer Data Lab (CSG), K01ES025434 award by NIEHS through funds provided by the trans-NIH Big Data to Knowledge (BD2K) initiative (LXG), P20 COBRE GM103457 award by NIH/NIGMS (LXG), R01 LM012373 award by NLM (LXG), R01 HD084633 award by NICHD (LXG), the Department of Defense BCRP [award number BC140682P1 (ACC)], the National Science and Engineering Council of Canada [NSERC DG grant number RGPIN-2016-05017 (AN)], the Windsor-Essex County Cancer Centre Foundation [Seeds4Hope grant number 814221 (AN)], Hopkins inHealth and Booz Allen Hamilton (90056858) to Y.X., the Russian Foundation for Basic Research KOMFI 17-00-00208 an NIH: NCI P30 CA006973 to A.V.F and National Research Council Canada to Y.F.. Views and opinions of, and endorsements by the author(s) do not reflect those of the US Army or the Department of Defense.

## Box 1: Technical description of matrix factorization

Matrix factorization approximates an input matrix x, as a product of two matrices, *U* and *V*, also referred to as factors.

In genomics, the input matrix *X* typically represents a preprocessed data matrix that has n molecular measurements as rows and p biological samples as columns (i.e., a matrix of size *n* x *p*). Often, not all of the row and column vectors of the data are equally informative. Exploratory data analysis aims to find a set of *k* most informative dimensions or features, wherekis smaller than either *n* or *p*. A *k*-dimensional representation that captures most of the “information” (say, for example, most of the variation) contained in the original data can then be used in its place in subsequent analysis.

MF solves this problem by approximating *X* as a product of two “factor” matrices: *X* ≈ *UV*, where the factor *U* is of size *n* x *k* and the factor *V* is of size *k* x *p*. Here, we refer to *U* as the amplitude matrix and *V* as the pattern matrix. We note that alternative variable names to *U* and *V* and terminology are often used in the literature, depending on the application.

Finding matrices *U* and *V* requires a quantitative definition of how well the product *UV* approximates *X*. One method for providing this definition is to use a loss function that quantifies the discrepancy between the approximation and the data, such as the mean square error and median absolute error. A common approach to solve for *U* and *V* is to apply an iterative approach such as a gradient-based method to minimize a loss function. Bayesian approaches to MF are alternatives to approximate the probabilistic relationship between *UV* and *X*. Additional conditions can also be incorporated in both gradient-based and Bayesian MF to learn useful features, such as sparsity constraints to limit the number of non-zero matrix elements.

Both gradient-based and Bayesian MF approaches have numerous applications beyond genomics. These techniques are ubiquitous in unsupervised feature learning for big data analysis.

## Box 2: PCA, ICA, and NMF

MF can start with the same data matrix *X* and then solve different problems to learn amplitude (*U*) and pattern (*V*) matrices with specific properties. An MF method requires: 1) the desired dimension *k*; 2) a loss function that measures the approximation *X* ≈ *UV*; and 3) constraints on u and/or v that enforce a desired structure on the low-dimensional representation. Changes to any of these give rises to different MF problems with different low-dimensional representations of the data. Three common MF approaches are applied to genomics: PCA, ICA, and NMF.

PCA finds a low-dimensional representation of the data that maximizes the variation contained in the original data. In PCA, the columns of uare orthogonal to each other describing non-overlapping structure in the data. Each column of represents the weight or loading of the data vector in the corresponding column of matrix *X*. In PCA, the columns of *U* and rows of *V* are ranked by the relative amount of variance they explain in the data. Thus, the first column of *U* and row of *V* explain most of the variation across molecular measurements and samples, respectively. Together, the *k* vectors learned with PCA maximize the total variance in the data captured using any rank *k* factorization. Geometrically, the vectors can be thought of as a set of orthogonal coordinate axes in high dimensional space, which represent the directions of maximal variation in the data.

ICA assumes that there are a set of *k* independent sources of variation that give rise to the observed data matrix *X*. This method enforces that the columns of *U* yields components that are statistically independent of each other. ICA is solved by minimizing the total mutual information between the *k* estimated components. The resulting factors ideally represent independent sources of variation in the biological system.

Non-negative matrix factorization (NMF) is a group of algorithms that constrains all elements of the *U* and *V* matrices to be greater than or equal to zero. The non-negativity constraint makes the representation purely additive, with no sources that can explain the data by removing signal. This non-negativity often results in NMF producing a *sparse* representation of the data. The additivity and sparsity make the *k* features inferred from NMF easy to interpret as the set of *active* components of the data that will give the original data when added together.

## Box 3: Common terminology in the literature

Historically, the independent discovery of MF in multiple fields including mathematics, computer science, and statistics created distinct terminologies that are often used interchangeably as analytical orthologs in genomics. For example, the term factorization is often used interchangeably with decomposition. Other terms, such as features, components, latent variable, or latent factors are both used to to refer to relationships between molecular measurements or samples, depending on the context.

The specific terminology for the amplitude and pattern matrices also varies by method and preferably labeled with different variable name. In PCA, the amplitude matrix *u* is often called the score or rotation matrix; and pattern matrix *V* is called the loadings. In ICA, the amplitude matrix is called the unmixing matrix (labeled as *A*) and the pattern matrix is called the source matrix (labeled as 5). In NMF, the amplitude matrix is commonly called the weights matrix (labeled as *W*) and the pattern matrix is called the features matrix (labeled as *H*).

Throughout this paper, we use the amplitude matrix to refer to matrix that contains vectors which represent relationships between molecular measurements. Other terms used in the literature for these molecular relationships include modules, meta-pathways, or signatures. Similarly, we use the pattern matrix to refer to the matrix that contains vectors which represent relationships between samples. Other literature terms for these sample-level relationships include patterns, metagenes, eigengenes, sources, or controlling factors.

## Glossary

Amplitude matrix: The matrix learned from MF that contains molecules in rows and factors in columns. Each column represents the relative contribution of the genes in a factor, which can be used to define a molecular signature for a CBP.
Complex biological process (CBP): The coregulation or coordinated effect of multiple molecular species resulting in one or more phenotypes examples can range from activation of multiple proteins in a single cellular signaling pathway to epistatic regulation of development.
Computational microdissection: A computational method to learn the composition of a heterogeneous sample, e.g., the cell types in a tissue sample.
Independent Component Analysis (ICA): A MF technique that learns statistically independent factors.
Matrix Factorization (MF): A technique to approximate a data matrix by the product of two matrices (see Box 1), one of which we call the amplitude matrix and the other the pattern matrix.
Non-negative Matrix Factorization (NMF): A MF technique for which all elements of the amplitude and pattern matrices are greater than or equal to zero.
Pattern matrix: The matrix learned from MF that contains factors in rows and samples in columns. Each row represents the relative contribution of the samples in a factor, which can be used to define the relative activity of CBPs in each sample.
Principal Component Analysis (PCA): A MF technique that learns orthogonal factors ordered by the relative amount of variation of the data that they explain.

